# Accurate CNV identification from only a few cells with low GC bias in a single-molecule sequencing platform

**DOI:** 10.1101/2020.01.21.908897

**Authors:** Liang Hu, Qunting Lin, Pingyuan Xie, Lidong Zeng, Lichun Liu, Mengxian Huang, Qin Yan, Meng Zhang, Ge Lin

## Abstract

A technical problem of characterizing copy number variation of several cells with next-generation sequencing is the whole genome amplification induced bias. The result of CNVs and mosaicism detection is affected by the GC bias. Here, we report a rapid non-WGA sample preparation strategy for a single-molecule sequencing platform GenoCare1600. This approach, combined with a single-molecule sequencing platform that avoids the use of WGA and bridge PCR processes, can provide higher reliability with its lower GC bias. By combining our optimized Tn5-based transposon insertion approach with GenoCare, we successfully detected CNVs as small as 1.29M and mosaicism as small as 20%, which is consistent with next-generation sequencing (NGS) data. Moreover, our GenoCare-TTI protocol showed less GC bias and less Mad of Diff. These results suggest that the optimized TTI approach, together with the GenoCare1600 sequencing platform, is a promising option for CNV characterization from maybe one single cell.

## INTRODUCTION

In vitro fertilization (IVF) is a technology that has been widely used in the treatment of infertility to improve pregnancy rates. However, a high proportion of first-trimester spontaneous miscarriages, which is associated with chromosomal aneuploidy in human pregnancies, greatly affected IVF outcome, especially for patients with advanced maternal age [1]. Preimplantation genetic testing for aneuploidy (PGT-A) can maximize the possibility of euploid embryo transfer, thus are thought to be a gospel to some IVF patients. Different genetic diagnostic technologies have been developed for PGT-A, such as fluorescence in situ hybridization (FISH) [2], quantitative polymerase chain reaction (qPCR) [3], microarray technologies including single nucleotide polymorphism (SNP) microarrays [4], array-based comparative genomic hybridization (aCGH) [5], and next-generation sequencing (NGS) [6]. Among them, FISH was not recommended for PGT-A since it could only screen a limited number of chromosomes, and the error rate of 5-15% usually led to disappointing pregnancy outcomes [7]. Array-based CGH and SNP microarray are reliable, but the expensive chip expenses significantly increase the PGT-A cost. The NGS approach is a widely used technique for PGT-A due to its ability to comprehensively screen all chromosomes at a competitive price. A recent study, using NGS platform, successfully detected CNV close to 1Mb in size [8]. Besides, NGS also can detect the presence of 20%-80% abnormal cells in a blastocyst biopsy [9]. The standard processes for PGT-A NGS library typically include whole genome amplification (WGA) of cells, fragmentation of WGA products, end-polishing, ligation of adaptor sequences, PCR amplification, and size-selection. However, WGA before NGS not only increases the PGT-A duration but also creates noises.

Recently, single-molecule sequencing arises as a new technology for clinical applications. It circumvents many library preparation issues by avoiding DNA amplification. In the past few years, previous work demonstrated sequencing of SNPs, M13 virus genome, and detection of trisomy 21/18/13 by single-molecule sequencing platform GenoCare [10 11]. Advantages of GenoCare include time-saving and straightforward sample preparation; (ii) absence of PCR amplification and low GC bias; (iii) significantly more sequencing reads than other single-molecule platforms.

WGA is a crucial step to enable comprehensive chromosome analysis. Over the years, there have been significant advances in WGA techniques, and several alternatives are available. The first method is primer extension pre-amplification (PEP), followed by the more widely adopted degenerate oligonucleotide-primed PCR (DOP-PCR). The basic principle of DOP-PCR is to use degenerate primers containing a random six-base sequence and a fixed sequence. However, in the process of PCR, due to uncertain factors such as input DNA amount and GC content, overamplified regions and unamplified regions appear, leading to amplification bias [12]. Multiple displacement amplification (MDA) was developed using isothermal amplification to solve this problem. In 2012, Zong et al. reported a single-cell WGA method called multiple annealing and looping-based amplicon cycles (MALBAC) that employed quasi-linear amplification through looping-based amplicon protection followed by PCR to reduce non-linear amplification [13]. Despite these advances in WGA for NGS-based PGT-A, Several issues still hinder its widespread clinical application: (1) Time: WGA and NGS library constructions require two days from beginning to the end with common workflows: WGA, DNA fragmentation and repair, ligation to specific adapters, PCR enrichment. (2) Cost: reagents and equipment used in this process are expensive. (3) Bias: Amplification biases are generated during the WGA and sequencing library construction [14]. The Tn5 transposase-based sequencing library preparation was then applied to address these problems. The enzyme catalyzes translocation by integrating the ME sequence into the target sites of DNA strands [15 16]. Due to its fast workflow, low DNA input, and limited hands-on time, this method has been used widely by researchers [17].

In this study, we developed a new library preparation method based on Tn5 transposase (**Figure 1**). Workflow and experimental conditions were optimized to skip the MDA process and give less hands-on time and GC bias. It was validated by comparison with traditional methods through sequencing on single-molecule sequencer GenoCare1600. Samples with copy number variations (CNVs) were also sequenced to demonstrate the ability to identify aneuploidy and mosaicism.

**Figure 1.**
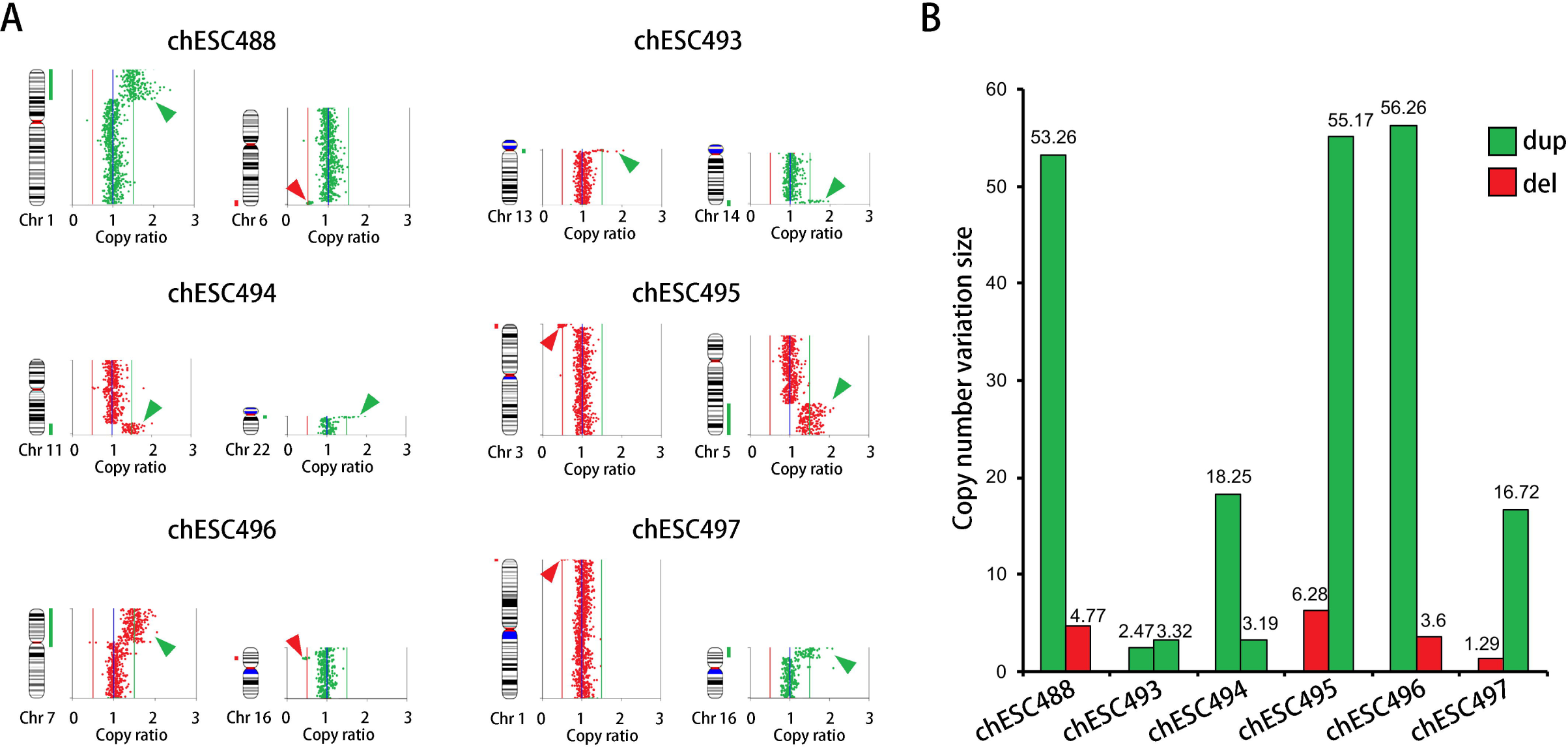
Next-generation sequencing (NGS) results of the six abnormal human embryonic stem cell lines. (A) The copy number variation (CNV) of affected chromosomes; (B) The CNV size of the six abnormal human embryonic stem cell lines.

## MATERIALS AND METHODS

### Cell lines

The cell lines used in this study were established and cultured in human embryonic stem cell (hESC) bank of the National Engineering Research Center of human Stem Cells [18]. In this paper, a 5-cell or 50-cell sample was picked up by microinjection. chHES90 was a normal hESC and was used to demonstrate the impact of MDA, and the other hESC lines were used for the comparison of library preparation methods and sequencing platforms.

### Tn5 transposon

To accommodate with single-molecule sequencer GenoCare, the adapter sequences for Tn5were designed as follows:

> ME-SEQ: 5’-[phos]-<u>CTGTCTCTTATACACATCT</u>-[NH_2_]-3’;
>
> Adapter 1:
>
> 5’-TCCTTGATACCTGCGACCATCCAGTTCCACTC<u>AGATGTGTATAAGAGACAG</u>-3’;
>
> Adapter 2:
>
> 5’-CTCAGATCCTACAACGACGCTCTACCGATGA<u>AGATGTGTATAAGAGACAG</u>-3’.

### Library construction via transposon insertion

Three library preparation methods were developed. In each case, 50ng gDNA was used as DNA input. Fragmentation and transposon insertion reactions were done as follows: 4μL 5×buffer L, 1μL target DNA at 50 ng/ μL, 1.5 μL Tn5 transposase, and 13.5 μL H_2_O were mixed and incubated at 55°C for 10min in a preheated thermocycler. After this mutual treatment, in route 1, DNA was purified by 1 volume VAHTS DNA clean bead (Vazyme), and 20μL solution was then transferred into a PCR tube, following by a five-cycle PCR process with 10 μL 5× GM PCR buffer, 2 μL forward primer, 2 μL reverse primer, 1 μL GM DNA polymerase, and 15 μL H_2_O. Afterward, the solution was purified by 1.2 volume VAHTS DNA clean beads, and the final solution was 20μL.

Route 2 is different from route 1 by adopting asymmetrical PCR amplification. The solution mixture contained 10 μL 5×GM PCR buffer, 2 μL forward primer, 0.2 μL reverse primer, 1 μL GM DNA polymerase, and 16.8 μL H_2_O. Ten cycles of PCR were performed to produce a sufficient yield. Eventually, a 20 μL solution was obtained after beads purification.

Route 3, after fragmentation and transposon insertion, the reaction was terminated by adding 5 μL 5× stop buffer and staying at room temperature for 5 min. No further purification was needed. Asymmetric PCR amplification was carried out for 10 cycles under the condition of 10 μL 5× GM PCR buffer, 2 μL forward primer, 0.2 μL reverse primer, 1 μL GM DNA polymerase, and 11.8 μL H_2_O. Eventually, a 20μL solution was obtained after beads purification.

From 50 ng DNA, we routinely got more than 25 ng/uL dsDNA from route 1 and 2, and ~200 ng/uL ssDNA from route 3. Products have a size distribution of 150~1000 bp and can be directly used on GenoCare.

Also, we develop a rapid library preparation kit base on Tn5 for cell lysis. In this paper, the GenoCare library was prepared according to the manufacturer’s protocol: gDNA was extracted from 5 or 50 cells. Six microliters of lysis buffer were added, spun down, and the DNA was incubated at 55°C for 1 h in a preheated thermocycler. 0.5 μL lysis stop buffer was then added, spun down, and the tube stayed at room temperature for 35 min to stop lysis reaction. No further purification was needed. The primers oligonucleotide sequences were as follows:

> forward primer 5’-TTCCTCAGATCCTACAACGACGCTCTACCGAT-3’
>
> reverse primer 5’-TTCTCCTTGATACCTGCGACCATCCAGTT-3’.

### MDA

MDA was performed on five human cells, as described in Vazyme Discover-sc single-cell kit (Vazyme). Briefly, 3μL cell lysis buffer was freshly prepared and added into each tube. After heating at 65°C for 10 min, 3 μL buffer N was added to stop the lysis reaction. 40 μL of amplification buffer was then added to start the MDA process. PCR steps were as follows: 30°C for 6 hours, 65°C for 3 min. In order to understand the relationship between amplification time and GC bias, 0.5h, 1h, 2h, 4h, and 6h reaction times were studied. The final products were characterized by Qubit (Invitrogen) and agarose gel electrophoresis.

MDA product was according to the manufacturer’s protocol for GenoCare library preparation, and the library construction is about 1.5 hours. Meanwhile, 6-hour MDA amplification products were performed using TruePrep DNA Library Prep Kit V2 for Illumina (Vazyme) and sequenced on Illumina Hiseq X10 as a comparison. The library preparation workflow of this NGS method is about 12 hours.

### Sequencing and data analysis

The samples were sequenced on Genemind Bioscience’s single-molecule sequencer GenoCare and Illumina’s HiseqX10 sequencer, yielding more than 4% genome coverage for each sample. GenoCare sequencing was performed according to the previously disclosed protocol [11]. For each sample, 25% of the area of each flowcell channel was imaged, and 72 cycles of sequencing data were collected, yielding more than 4 million reads. NGS Sequencing was done on Illumina Hiseq X10 with standard PE-150 protocol according to the operating instruction manual, and the total sequence time is about three days.

In a basic data analysis process, Illumina X10 sequencing data were mapped to the reference human genome (hg19) by bwa, and antigenocide data were mapped to the same reference using home-written software called DirectAlign. Raw reads with low quality and non-unique alignments were removed. To reduce the influence of GC and mappability differences, we split the reference into 150 Kbp windows and kept the bins with GC content 32%~60% and mappability bigger than 0.6. To compare GC bias between samples, we calculated the relative bin density (RBD) by *R_i,j_=r_i,j_/M*. Where r denotes reads number in each bin, *i* and *j* represent different bins and samples, and M is the average number of sequencing reads in bins on autosomes. GC bias (Δ*R_GC_^2^*) was defined as

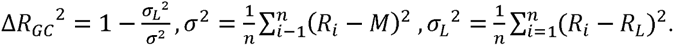

Moreover, *R_L_* represents the optimal prediction, which was obtained via a loess regression fit of the RBD against the GC content. Then we developed a weighted correction strategy to correct GC content and mappability in the scale of every 0.1% bin. Weight index *w* was calculated from an R function loss; the corrected bin density (CBD) was calculated in the following formula: *CBD=RBD/w*. This normalized bin density is used for further analysis. CNV identification was made by R packages (DNA copy).

We calculated the median absolute deviation of difference (Mad of Diff) to evaluate the reproducibility instead of the coefficient of variation (CV) because Mad of Diff represents the difference of corrected and normalized adjacent bin copy number, which is more accurate when assessing CNV samples.

### Statistics

The differences in coverage, GC bias, and Mad of Diff were compared using the student’s t-test. P<0.05 was considered statistically significant. Analyses were performed using the statistical package SPSS, version 18.0 (SPSS).

## RESULTS

### Cell line

To develop an effective library method strategy for CNVs and mosaicism detection at the cell level through a single-molecule sequencing platform, we chose five human embryonic stem cell lines with different sizes of CNV range from 1.29 Mbp to 56.26 Mbp (**Table 1**). NGS confirmed all the CNVs of the cells lines with their total genomic DNA (**Figure 1A**). The range of copy number gain (CNG), whose copy ratios were 3/2, were from 2.47 Mbp to 56.26 Mbp; And the range of copy number loss (CNL), whose copy ratios were 1/2, were from 1.29 Mbp to 6.28 Mbp (**Figure 1B**).

**Table 1.** Cell lines information

| Cell line name | CNV-Seq result |
| --- | --- |
| chHES488 | 46, XY, dup (2p25.3-2p16.2) 53.26 M, del (6q27) 4.77 M |
| chHES493 | 46, XY, dup (13q12.11) 2.47 M, dup (14q32.31-14q32.33) 3.32 M |
| chHES494 | 46, XX, dup (11q23.3-11q25) (18.25 M), dup (22q11.1-22q11.21) 3.19 M |
| chHES495 | 46, XX, del (3p26.3-3p26.1) 6.28 M, dup (5q23.2-5q35.3) 55.17 M |
| chHES496 | 46, XX, dup (7p22.3-7p11.2) 56.26 M, del (16p13.11-16p12.3) 3.60 M |
| chHES497 | 46, XX, del (1p36.33-1p36.32) (1.29 M), dup (16p13.3-16.13.11) 16.72 M |
| chHES90 | 46, XY |

### Whole-genome amplification induced significant bias

It is known that different WGA approaches induce error and template bias. To evaluate the influence of short-time MDA, we sequenced the short-time MDA products (from 0.5 hours to 6 hours) of five chESC90 cells and compared with those of unamplified 5-cell DNA, 50-cell DNA, and DNA of bulk cells (**Figure 2A**). To minimize the effect of amplification, we used Tn5-based transposon insertion (TTI) to construct the library for next-generation sequencing. As expected, the coverage of unamplified 5-cell DNA and 50-cell DNA was lower than that of bulk DNA and MDA-amplified products (**Figure 2B**). Surprisingly, even MDA for as short as 30 minutes induced statistically significant bias on GC bias and Mad of Diff. As shown in Figure 2B, after an only 30-minute amplification, the GC bias increased to 0.2. After 6-hour MDA, GC bias even reached as high as 0.62, which was 62 fold compare to bulk DNA without amplification (0.01).

**Figure 2.**
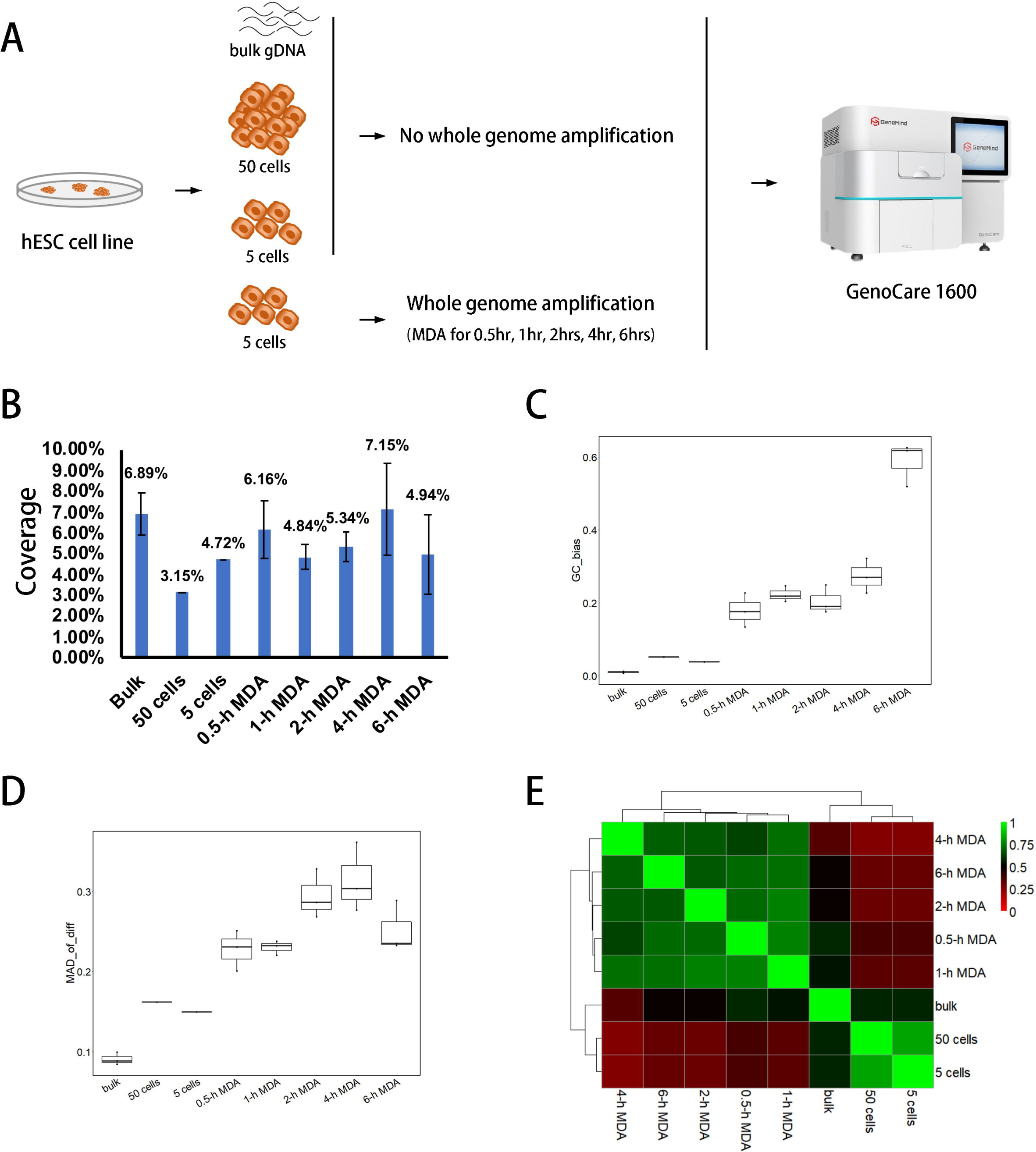
WGA induced significant bias in NGS. (A) The experimental design scheme to characterize the WGA-induced bias; (B) Unamplified gDNA had relatively low coverage; (C) Unamplified gDNA had less GC bias; (D) Unamplified gDNA had less Mad of Diff; (E) Amplified gDNA and unamplified gDNA correlated together, respectively.

Importantly, although the GC bias of unamplified 5 cells and 50 cells also increased to 0.03 and 0.05 respectively, they were only about 1/5 of that of 0.5-hour MDA, indicating that cell number does not significantly affect GC bias (**Figure 2C**). Similarly, Mad of Diff, a criterion which reflects the standard deviation of the sequencing data, increased to about 3 fold of that of bulk DNA (0.09) after amplification. Moreover, the Mad of Diff only increased to 0.15 for unamplified 5 cells and 50 cells (**Figure 2D**). The results were summarized in **Table S1**.

To get further insight into correlations between those library preparation methods, we calculated Pearson’s cross-correlation coefficients of relative bin density (RBD) between each sequencing result (**Figure 2E**). The hierarchical clustering of the correlation coefficient matrix showed that the MDA methods were well separated from each other and were also very different than the PCR-free bulk cells. Results from unamplified 5 cells and 50 cells and bulk cells are highly correlated, indicating that cell lysis following by unamplified Tn5-based library preparation had the least bias.

### Optimization of Tn5-based transposon insertion protocol

Our above data showed that Tn5-based transposon insertion (TTI) is a promising library prep strategy since the DNA was unamplified. However, amplification of the trace amount of DNA template is necessary during PGT-A, especially when the patients choose both PGT-A and PGT for monogenic diseases (PGT-M). Thus, we developed three routes of different pretreatment protocols and PCR strategies to optimize the amplification-based TTI (**Figure 3A**). Route 1 is a frequently used approach that purifies the DNA by beads and then amplified the purified DNA by symmetrical PCR. Since symmetrical PCR might lose some of single-strand DNA (ssDNA), we amplified the beads-purified DNA by asymmetrical PCR (Route 2). To minimize the loss of DNA during purification, we just use stop buffer to stop the TTI reaction and then amplified the unpurified DNA by asymmetrical PCR (Route 3). To test the performance of three routes, we sequenced six hESC cell line MDA product samples on the GenoCare platform and analyzed sequencing results of unique data, coverage, GC bias, and Mad of Diff (**Table S2**).

**Figure 3.**
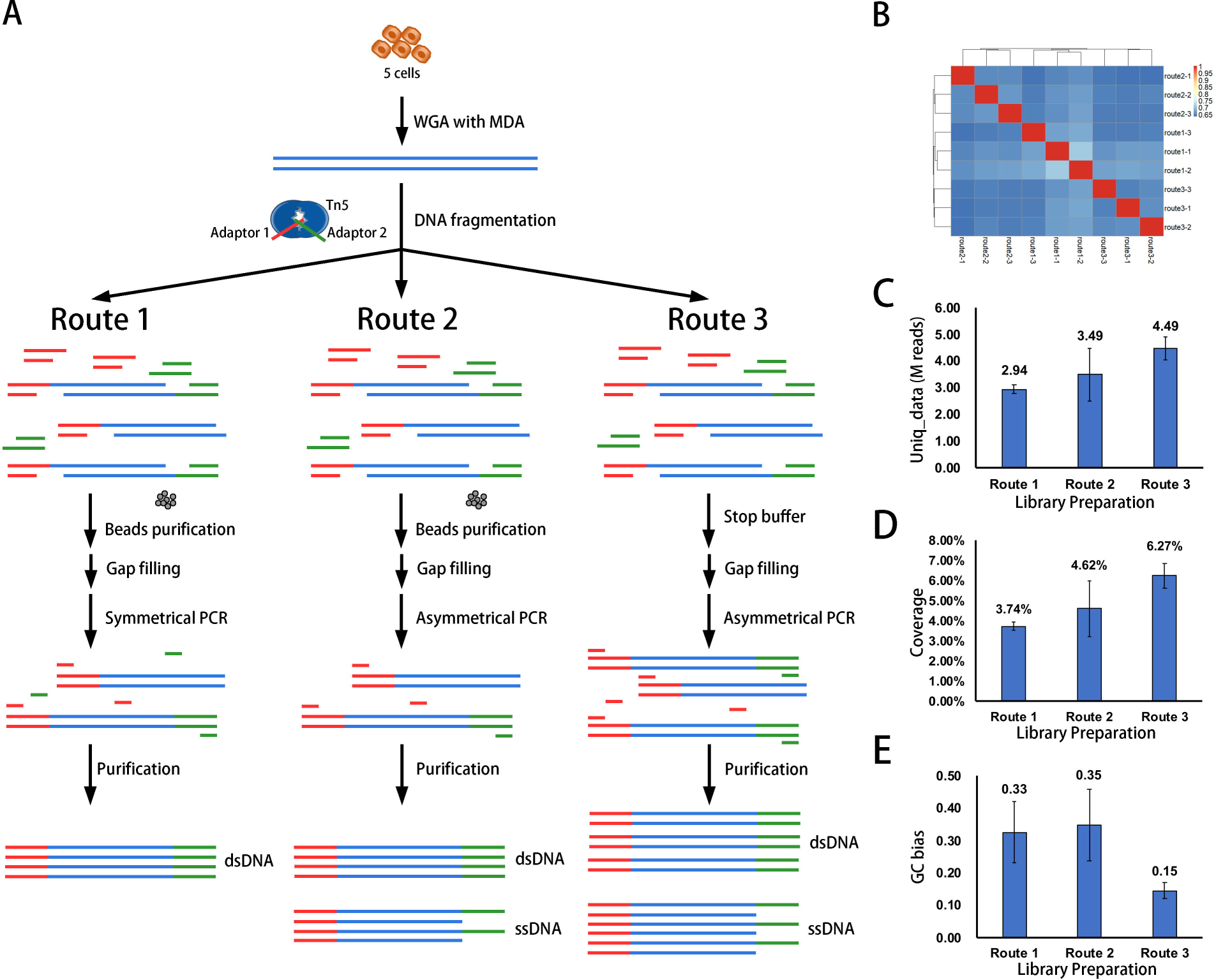
Stop buffer, and asymmetrical PCR optimized the Tn5-based transposon insertion (TTI) protocol. (A) The experimental design scheme to optimize the TTI protocol; (B) The sequencing results of three routes had a weak correlation with each other; (C, D, E) Route 3 had more unique data, higher coverage, and less GC bias.

We first calculated the correlation coefficient matrix among the three routes. As shown in **Figure 3B**, the hierarchical clustering of the correlation coefficient matrix showed that different library preparation routes and conditions clustered, respectively. They also displayed a weak correlation with each other, which indicated that each route had its built-in pattern of the library.

As shown in **Figure 3C and 3D**, compared with other routes, route 3 can obtain the most unique data (route 3 vs. route 1, 4.49 vs. 2.94 M reads, p<0.01; route 3 vs. route 2, 4.49 vs. 3.94 M reads, p<0.05) and the highest coverage (route 3 vs. route 1, 6.27% vs. 3.74%, p<0.01; route 3 vs. route 2, 6.27 vs. 4.62%, p<0.05), indicating that the stop buffer and asymmetric PCR amplification can avoid the DNA loss. Besides, the average GC bias of route 3 is much less than the other two routes (**Figure 3E**, route 3 vs. route 1, 0.15 vs. 0.33, p<0.01; route 3 vs. route 2, 0.15 vs. 0.35, p<0.01).

Besides, since route 3 does not need the beads purification, its library preparation time is only 1.5 hours, which saves 0.5-1 hour compare to route 1 and route 2 (**Table S2**). The above data indicated that route 3 has a clear advantage over the other two routes, which makes it an ideal method for CNV detection for trace amount cells.

### Optimized TTI protocol has less GC bias and can identify small CNVs

To test the performance of the optimized TTI protocol in identifying CNVs, we sequenced six hESC cell line gDNA samples with small CNVs on the GenoCare platform. The results were compared with those amplified by MDA and sequenced on Illumina HiseqX10 with 150bp paired-end reads. As shown in **Table 2**, the average coverage of GenoCare-TTI protocol was less than that of X10-MDA protocol (7.61% vs. 14.90), because the average read length of GenoCare1600 was less than that of Illumina HiseqX10 (42.06 vs. 150.00). Notably, the GC bias of GenoCare-TTI protocol was much less than that of the X10-MDA protocol (0.03 vs. 0.23, p<0.01), indicating that GenoCare-TTI protocol might have better coverage with fewer reads. Moreover, the average Mad of Diff of GenoCare-TTI protocol was less than that of X10-MDA protocol (0.22 vs. 0.31, p<0.05), indicating that GenoCare-TTI protocol might be more powerful to detect small CNVs.

**Table 2.** Comparison of different library preparation protocols and sequencing platforms

| Sample name | Platforms | Average<br>Uniq_ data<br>(M reads) | Average<br>Read length<br>(bp) | Average<br>coverage<br>(%) | Average<br>GC bias | Average<br>Mad of<br>Diff |
| --- | --- | --- | --- | --- | --- | --- |
| chHES493-MDA | Illumina Hiseq X10 | 4.77 | 150.00 | 14.62 | 0.32 | 0.23 |
| chHES495-MDA | Illumina Hiseq X10 | 5.72 | 150.00 | 16.77 | 0.21 | 0.35 |
| chHES497-MDA | Illumina Hiseq X10 | 4.97 | 150.00 | 13.31 | 0.15 | 0.34 |
| **MDA** | **Illumina Hiseq X10** | **5.15** | **150.00** | **14.90** | **0.23** | **0.31** |
| chHES493-TTI | GenoCare 1600 | 5.72 | 42.14 | 8.03 | 0.02 | 0.23 |
| chHES495-TTI | GenoCare 1600 | 5.22 | 41.93 | 7.30 | 0.03 | 0.24 |
| chHES497-TTI | GenoCare 1600 | 5.35 | 42.10 | 7.51 | 0.04 | 0.20 |
| **TTI** | **GenoCare 1600** | **5.43** | **42.06** | **7.61** | **0.03** | **0.22** |

Besides, we also evaluated whether GenoCare-TTI protocol can identify the small CNVs. As shown in **Figure 4A**, all the CNVs, including a 1.29M small CNV, can be successfully identified both with GenoCare-TTI protocol and with X10-MDA protocol. It should be noted that since GenoCare-TTI protocol has less Mad of Diff, the small CNV was more easily to be characterized than X10-MDA protocol (**Figure 4B**).

**Figure 4.**
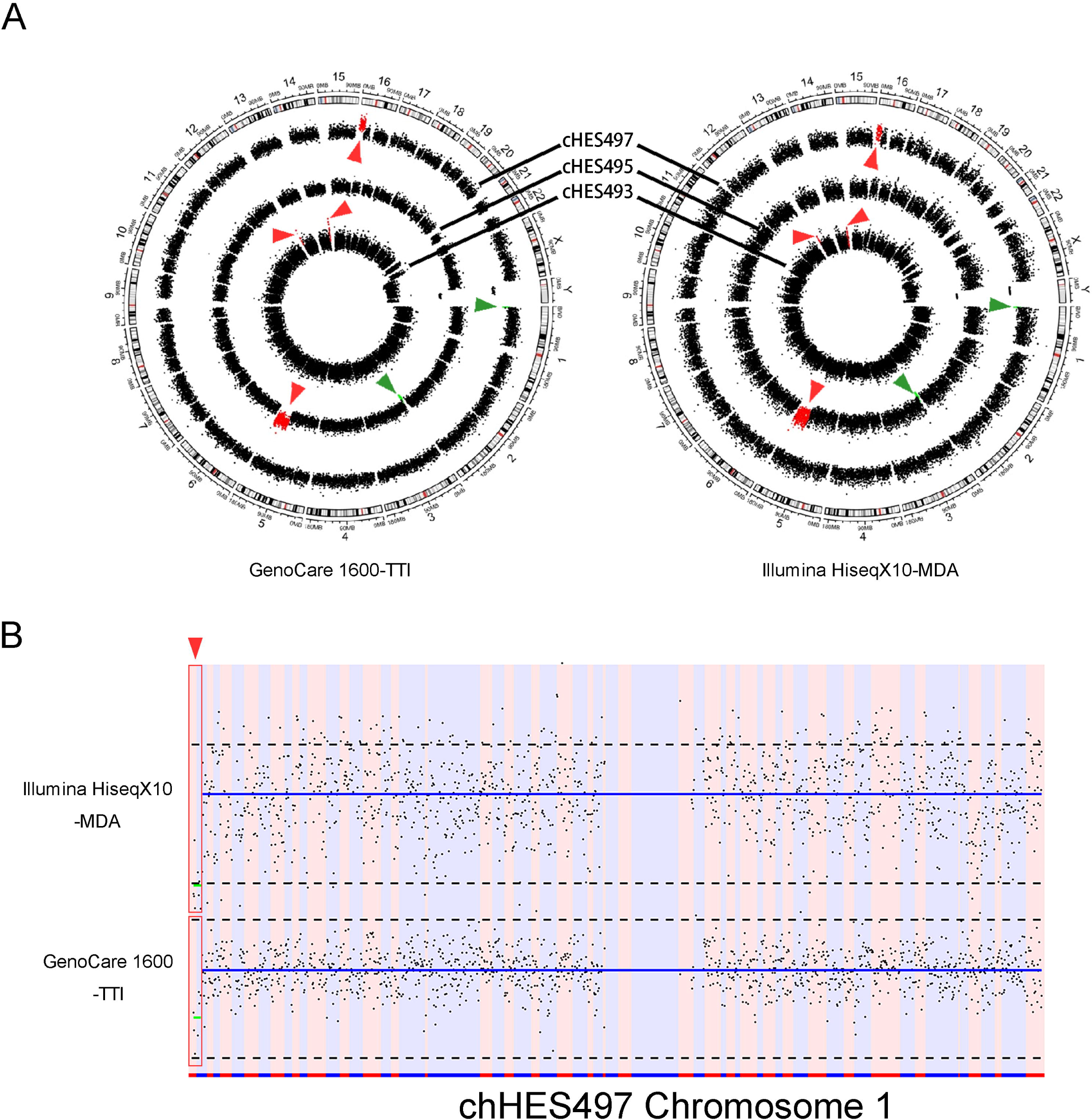
Optimized TTI protocol can identify small CNVs. (A) The small CNVs can be identified by both GenoCare-TTI protocol and X10-MDA protocol; (B) Both protocols could detect a CNV as small as 1.29M. Moreover, X10-MDA protocol had less Mad of Diff.

### Optimized TTI protocol can identify mosaicism

To investigate the accuracy of PGT-A in mosaicism detection, we made a series of mosaicism samples by mixing CNV-cell line chHES488 with the normal hESC cell line 90-P51 (**Figure 5A**). Those mosaic samples were characterized by either GenoCare-TTI protocol or with X10-MDA protocol. As shown in **Figure 5B**, even 20% of mosaicism can be identified by both protocols. We also found that GenoCare-TTI protocol has much less GC bias (**Table S3**) in this experiment, the average bin copy number remained stable as the GC content varied (**Figure 5C**).

**Figure 5.**
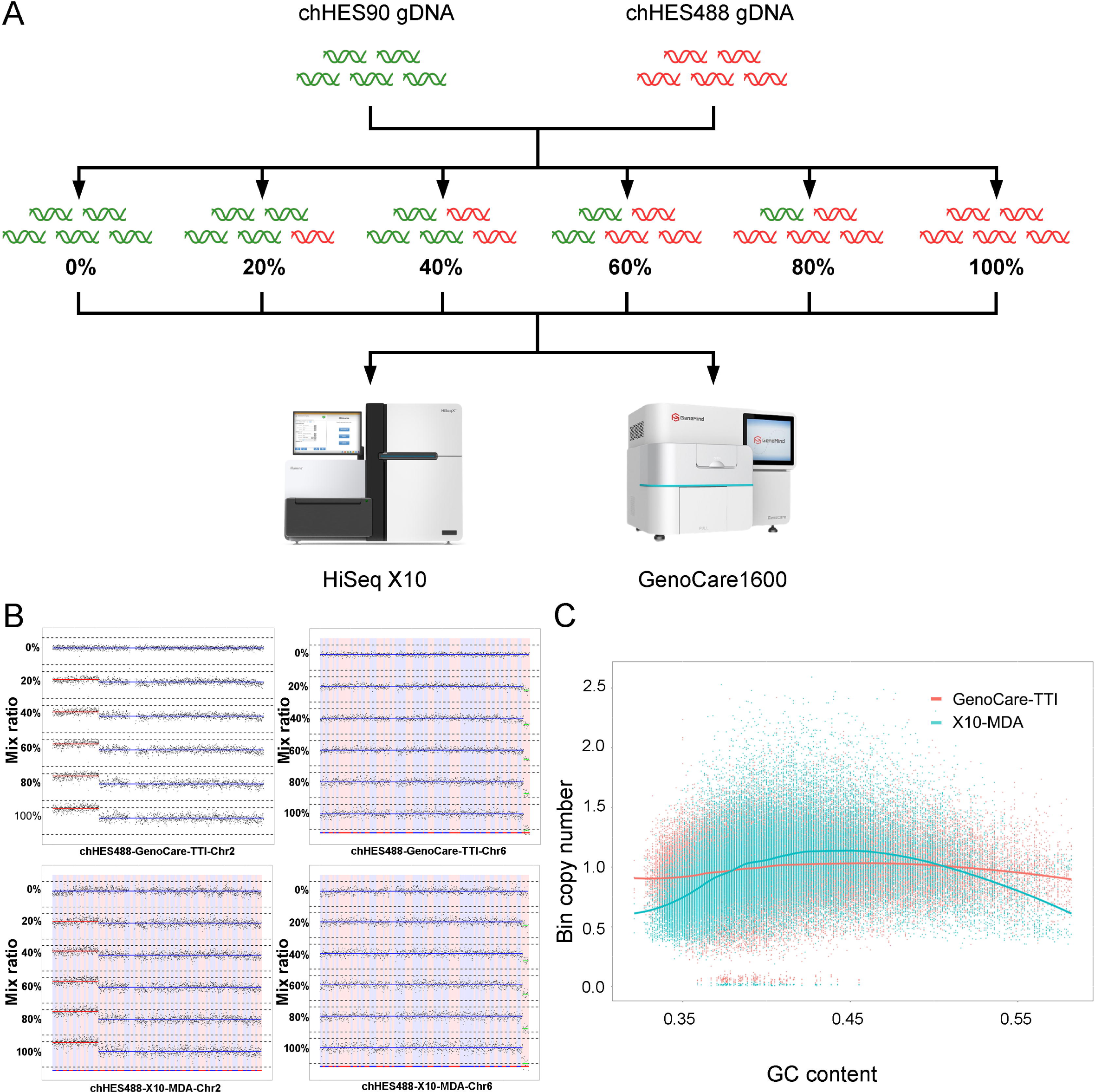
Optimized TTI protocol can identify as low as 20% mosaicism. (A) The experimental design scheme of mosaicism characterization; (B) Both protocols could detect as low as 20% mosaicism; (C) The average bin copy number of GenoCare-TTI protocol did not vary with GC content. The bin coverage rate distribution across the genome from different libraries preparation and a bulk-cell sample. Each bar represents the coverage rate in a 150 kb bin, and all 17019 bins are plotted.

## DISCUSSION

Our present study demonstrated the validity of a new 1.5-hour PGT-A sample preparation method for as little as five cells, like the blastocyst biopsy samples during PGT-A. The technical accuracy was measured in MDA amplification products CNVs and mosaic ratios. We compared the influence of the gain achieved in MDA on the characteristics of interest. Overall, the amplification bias in MDA is a direct function of the overall reaction gain, with more significant gain leading to more bias. This observation underlines the importance of tailoring the gain of amplification to yield a higher quality of DNA products for the subsequent sequencing workflow. The overall gain can be set through by reducing the reaction time.

For single-cell sequencing, WGA is required before sequencing library construction during NGS based PGT-A. It has been demonstrated that significant GC bias will arise during the WGA process, which would affect the sequencing profile accuracy, leading to false positive or false negative [19]. In our study, we found that the cell genome amplification by cell lysis Tn5-based library preparation method is highly efficient. It rendered better reproducibility and uniformity than MDA, especially concerning GC content.

We also compared the efficiency of CNVs detection by different Tn5-based library preparation methods. All three optimized Tn5-based library preparation methods generated sequence data on the SMTS GenoCare platform and are concordant with the date from the Illumina X10 platform. Among the three routes, Route 1 and 2 showed high GC bias, which may be caused by two reasons. Previous research revealed that half of the genomic DNA fragments were lost due to transposon symmetry in the conventional method [20]. As a result, PCR amplification lost uniformity due to the lower template amount.

Moreover, since the same adaptor sequences were attached to genomic DNA fragments on both ends, self-looping of the template strand may happen, which could lead to PCR amplification failure in the GC-rich regions. To solve this problem, we designed an asymmetric PCR amplification to enrich ssDNA [21]. Meanwhile, we replaced beads purification with stop buffer to stop fragmentation reaction to reduce the operation time and DNA loss. Interestingly, we found that a combination of asymmetric PCR and stop buffer treatment (route 3) could not only yield reliable results for CNVs diagnosis but also was faster than the other two Tn5-based library preparation methods (route 1 and 2). Meanwhile, NGS results of route 3 showed much less GC bias with compare to those of route 1 and 2. Therefore, route 3 may be an ideal method for PGT-A.

Mosaicism in embryos as a result of postzygotic mitotic aneuploidy could contribute to biologic variation in blastocysts. Chromosomal mosaicism frequently occurs in human blastocysts as detected during PGT-A [22]. A mosaic embryo or cell line is that one has cells with different CNVs in at least one chromosome. Levels of mosaicism at the cleavage stage were estimated to range from 15% to 90% [23]. According to the Preimplantation Genetic Diagnosis International Society (PGDIS), less than 20% mosaicism is deemed as euploidy. When the mosaic ratio is more than 80%, the embryo is considered to be aneuploidy. When the mosaic ratio is between 20% and 80%, PGT-A results will help doctors make embryo transfer decisions. Empirically, >50% mosaic embryos are easy to detect with the use of multiple PGT-A platforms. To fully validate route 3 for PGT-A, we also applied it on the mosaic curve detection. As our data showed, the six different proportions of mosaic samples of this curve have a noticeable trend of gradually decreasing. Analysis of these well-controlled samples by route 3 demonstrated perfect consistency with the expected mixing proportions. The standard curve of the mosaic ratio can provide a relatively accurate reference method for the assessment of the chimeric ratio during PGT-A. The approach may help reduce the impact of mosaicism and biologic variation on evaluating the technical accuracy of new methods. Route 3, combining with SMTS GenoCare 1600 sequencer, can accurately and reproducibly measure 20% mosaicism in a known sample.

To our knowledge, there were very few reports that a single molecule sequencer was applied in PGT-A. In this study, we compared the sequencing performance of GenoCare1600 and Illumina Hiseq X10 for as little as five cells in the same 11 hESC lines to mimic the real PGT-A circumstances. At least 4 million sequencing reads were collected for each sample, which meets the requirement of PGDIS. With the PE-150 sequencing protocol, X10 delivered higher RL and genome coverage, while GenoCare 1600 gave lower GC bias and shorter sequencing time with less sequencing cycles. GenoCare 1600 could produce more than 10 million reads per flowcell channel, 160 million reads per flowcell, comfortably accommodating 16 PGT-A samples for each run. Only one-fourth of each flowcell channel was imaged in this study to reduce sequencing time further. The reads number is one to two orders of magnitude higher than other single-molecule sequencers, namely Pacific biosciences Sequeland Oxford Nanopore’s Minion/GridIon. Although GenoCare’s read length is short, its large amount of reads number makes it more suitable for CNV detection.

In this study, we characterized an alternative method of library construction, which combines with Genocare sequencing platform produced fast and accurate PGT-A data. We have successfully used the Tn5-based library preparation method to investigate the different CNVs of the cell-line samples, which was consistent with NGS results. Besides, for samples with a 20% to 80% mosaic ratio, Genocare sequencing results are very consistent with NGS results. We believe that this new platform delivers a competitive solution to PGT-A by reducing the test time and generating data with lower GC bias and comparable CNV sensitivity.

## ACKNOWLEDGMENTS

Supported by the National Key R&D Program of China (grant 2018YFC1003100, to L.H.), the National Natural Science Foundation of China (grant 81873478, to L.H.), and the science and technology major project of the ministry of science and technology of Hunan Province, China (grant 2017SK1030, to G.L.).

