## Supplementary material for "Accurate CNV identification from only a few cells with low GC bias in a single-molecule sequencing platform": Table S1

Table S1. Whole genome amplification induced significant bias

| Sample name | Uniq_data (M reads) | Read length (bp) | coverage | GC bias | Mad of Diff |
| --- | --- | --- | --- | --- | --- |
| Bulk | 5.62 | 37.67 | 6.89% | 0.01 | 0.09 |
| 50 cells | 2.63 | 36.11 | 3.15% | 0.05 | 0.16 |
| 5 cells | 4.13 | 35.31 | 4.72% | 0.03 | 0.15 |
| 0.5-h MDA | 5.08 | 36.98 | 6.16% | 0.18 | 0.23 |
| 1-h MDA | 4.15 | 35.30 | 4.84% | 0.22 | 0.23 |
| 2-h MDA | 4.55 | 35.68 | 5.34% | 0.21 | 0.30 |
| 4-h MDA | 5.64 | 39.29 | 7.15% | 0.27 | 0.31 |
| 6-h MDA | 3.85 | 38.88 | 4.94% | 0.59 | 0.25 |
