## Supplementary material for "Accurate CNV identification from only a few cells with low GC bias in a single-molecule sequencing platform": Table S2

Table 2. The validation of different library methods

| Sample name | Library Prep | operation time (h) | Uniq_data | Read length (bp) | coverage | GC bias | Mad of Diff |
| --- | --- | --- | --- | --- | --- | --- | --- |
| chHES488 | Route 1 | 2.0 | 3040464 | 38.37 | 3.89% | 0.37 | 0.15 |
|  | Route 2 | 2.5 | 4202462 | 40.35 | 5.65% | 0.48 | 0.14 |
|  | Route 3 | 1.5 | 5107057 | 41.87 | 7.13% | 0.13 | 0.14 |
| chHES493 | Route 1 | 2.0 | 3115529 | 38.30 | 3.98% | 0.39 | 0.23 |
|  | Route 2 | 2.5 | 1752497 | 38.95 | 2.28% | 0.36 | 0.25 |
|  | Route 3 | 1.5 | 4290289 | 41.90 | 5.99% | 0.16 | 0.22 |
| chHES494 | Route 1 | 2.0 | 3055375 | 38.20 | 3.89% | 0.46 | 0.27 |
|  | Route 2 | 2.5 | 4084491 | 40.51 | 5.52% | 0.48 | 0.27 |
|  | Route 3 | 1.5 | 4064206 | 41.71 | 5.65% | 0.15 | 0.27 |
| chHES495 | Route 1 | 2.0 | 2911606 | 38.21 | 3.71% | 0.25 | 0.33 |
|  | Route 2 | 2.5 h | 4413724 | 40.26 | 5.92% | 0.29 | 0.32 |
|  | Route 3 | 1.5 h | 4041800 | 41.56 | 5.60% | 0.11 | 0.33 |
| chHES496 | Route 1 | 2 h | 2721733 | 38.12 | 3.46% | 0.24 | 0.34 |
|  | Route 2 | 2.5 h | 3051606 | 38.67 | 3.93% | 0.25 | 0.34 |
|  | Route 3 | 1.5 h | 4810690 | 41.90 | 6.72% | 0.14 | 0.34 |
| chHES497 | Route 1 | 2 h | 2788213 | 37.93 | 3.53% | 0.24 | 0.32 |
|  | Route 2 | 2.5 h | 3458810 | 38.38 | 4.43% | 0.23 | 0.32 |
|  | Route 3 | 1.5 h | 4639403 | 42.03 | 6.50% | 0.18 | 0.32 |
