## Supplementary material for "Accurate CNV identification from only a few cells with low GC bias in a single-molecule sequencing platform": Table S3

Table S3. GenoCare-TTI and X10-MDA sequencing results for mosaicism

| Protocol | Sample name | Uniq_ data (M reads) | Read length (bp) | coverage | GC bias | Mad of Diff |
| --- | --- | --- | --- | --- | --- | --- |
| GenoCare-TTI | 100%chHES488 | 5107057 | 41.87 | 7.13% | 0.13 | 0.14 |
|  | 80%chHES488 | 4184627 | 42.08 | 5.87% | 0.23 | 0.13 |
|  | 60%chHES488 | 4761372 | 42.08 | 6.68% | 0.27 | 0.12 |
|  | 40%chHES488 | 5180778 | 42.12 | 7.27% | 0.16 | 0.11 |
|  | 20%chHES488 | 5292530 | 42.30 | 7.46% | 0.04 | 0.10 |
|  | 0%chHES488 | 491000 | 41.50 | 6.79% | 0.029 | 0.08 |
| X10-MDA | 100%chHES488 | 4696221 | 150.00 | 15.08% | 0.35 | 0.17 |
|  | 80%chHES488 | 7900741 | 150.00 | 21.05% | 0.39 | 0.14 |
|  | 60%chHES488 | 7365912 | 150.00 | 17.94% | 0.26 | 0.13 |
|  | 40%chHES488 | 7480599 | 150.00 | 17.19% | 0.30 | 0.12 |
|  | 20%chHES488 | 7061456 | 150.00 | 16.48% | 0.52 | 0.12 |
|  | 0%chHES488 | 23953218 | 150.00 | 59.25% | 0.59 | 0.11 |
